# Neural activity during story listening is synchronized across individuals despite acoustic masking

**DOI:** 10.1101/2021.03.25.437022

**Authors:** Vanessa C. Irsik, Ingrid S. Johnsrude, Björn Herrmann

**Affiliations:** Department of Psychology & The Brain and Mind Institute, The University of Western Ontario, London, ON, Canada N6A 3K7; School of Communication and Speech Disorders, The University of Western Ontario, London, ON, N6A 5B7, Canada; Rotman Research Institute, Baycrest, M6A 2E1, Toronto, ON, Canada; Department of Psychology, University of Toronto, M5S 1A1, Toronto, ON, Canada

**Author notes:** Correspondence concerning this article should be addressed to Vanessa C. Irsik, The Brain and Mind Institute, The University of Western Ontario, London, Ontario, N6A 5B7, Canada.

**Keywords:** naturalistic stimuli, hearing, speech intelligibility, engagement, inter-subject correlation

## Abstract

Older people with hearing problems often experience difficulties understanding speech in the presence of background sound. As a result, they may disengage in social situations, which has been associated with negative psychosocial health outcomes. Measuring listening (dis-)engagement during challenging listening situations has received little attention thus far. We recruit young, normal-hearing human adults (both sexes) and investigate how speech intelligibility and engagement during naturalistic story listening is affected by the level of acoustic masking (12-talker babble) at different signal-to-noise ratios (SNRs). In Experiment 1, we observed that word-report scores were above 80% for all but the lowest SNR (-3 dB SNR) we tested, at which performance dropped to 54%. In Experiment 2, we calculated inter-subject correlation (ISC) using electroencephalography (EEG) data to identify dynamic spatial patterns of shared neural activity evoked by the stories. ISC has previously been used as a neural measure of participants’ engagement with naturalistic materials. Our results show that ISC was stable across all but the lowest SNRs, despite reduced speech intelligibility. Comparing ISC and intelligibility demonstrated that word-report performance declined more strongly with decreasing SNR compared to ISC. Our measure of neural engagement suggests that individuals remain engaged in story listening despite missing words due to background noise. Our work provides a potentially fruitful approach to investigate listener engagement with naturalistic, spoken stories that may be used to investigate (dis)engagement in older adults with hearing impairment.

## Introduction

Many listening environments we encounter contain background masking sounds (Olsen, 1998; Smeds et al., 2015) that can lead to listening challenges for a significant proportion of people aged 50 and older (Frisina & Frisina, 1997; Gordon-Salant, 2006). People who frequently experience listening difficulties may avoid environments with increased levels of background noise, resulting in social isolation (Dawes et al., 2015; Palmer et al., 2016) and negative psychosocial and physical health outcomes (Pichora-Fuller et al., 2015; Wayne & Johnsrude, 2015). However, social isolation is likely subsequent to within-situation disengagement; a coping mechanism to reduce cognitive demand. A person may temporarily “zone out” in conversational situations because continuous listening is too difficult (Heffernan et al., 2016; Herrmann and Johnsrude, 2020a, 2020b). A greater understanding of the listening conditions under which a person may disengage, and development of quantitative measures of within-situation disengagement, may improve our ability to diagnose hearing problems before social isolation manifests.

Two factors are likely critical for listening engagement, which we define as “the (automatic or volitional) recruitment of executive and other cognitive resources, when speech comprehension serves a valued communication goal” (Herrmann & Johnsrude, 2020a). First, engagement depends on degree to which a listener judges that, with deliberate deployment of cognitive resources, some of the masked speech will be understandable (Brehm & Self, 1989; Herrmann & Johnsrude, 2020a; Pichora-Fuller et al., 2016; Richter, 2013; Wright, 2014). An individual may disengage if they believe speech comprehension will be impossible (Eckert et al., 2016; Peelle, 2018; Richter, 2016). A second important factor is motivation. An individual who is motivated to listen, for example, because they find what they are hearing to be enjoyable and rewarding (Matthen, 2016), may engage in listening despite the presence of background sounds (Herrmann & Johnsrude, 2020a; Pichora-Fuller et al., 2016). However, typical investigations of speech comprehension involve listening to isolated sentences (Davis & Johnsrude, 2003; Duncan & Aarts, 2006; Obleser et al., 2007) that lack a topical thread, are not very interesting (e.g., He buttoned his shirt), and may not therefore foster volition to listen (i.e., conation; Reitan and Wolfson, 2000). Further, listening engagement also likely develops more slowly, at time scales beyond individual sentences (Mandler & Goodman, 1982). Spoken stories, in contrast, involve event descriptions along a topical thread, are intrinsically motivating to a listener, and are common in everyday life (Dunlop & Walker, 2013). Utilizing engaging spoken stories under varying levels of acoustic masking may thus provide a fruitful avenue to investigate intelligibility, and listening engagement and disengagement.

Engagement (and disengagement) can be measured behaviorally through assessment of an individual’s experience (Busselle & Bilandzic, 2008, 2009; Dmochowski et al., 2014; Herrmann & Johnsrude, 2020b; Kuijpers et al., 2014). However, behavioral assessment of engagement typically requires the listener to retroactively introspect about their experiences (Herrmann & Johnsrude, 2020b). In contrast, neural activity recorded using functional imaging or electroencephalography (EEG) may provide a real-time window on engagement. Naturalistic materials such as movies or spoken stories lead to synchronized patterns of neural activity across individuals that scale with the degree of engagement (Hasson et al., 2010; Nastase et al., 2019; Nguyen et al., 2019; Yeshurun et al., 2017). The strength of synchrony across individuals, quantified as inter-subject correlation (ISC; Hasson et al., 2004, 2008; Dmochowski et al., 2012, 2014), is stronger when stimuli are captivating or exciting (Hasson et al., 2010; Schmälzle et al., 2015), and is predictive of behavioral measures reflecting engagement (S. S. Cohen et al., 2017; Dikker et al., 2017; Dmochowski et al., 2014; Poulsen et al., 2017; Song et al., 2021) and recall of the materials (Chan et al., 2019; S. S. Cohen et al., 2018; S. S. Cohen & Parra, 2016; Davidesco et al., 2019; Hasson, Furman, et al., 2008; Piazza et al., 2021; Song et al., 2021; Stephens et al., 2010). Conversely, ISC is reduced when individuals do not attend to naturalistic materials (S. S. Cohen et al., 2018; Ki et al., 2016; Kuhlen et al., 2012; Rosenkranz et al., 2021) or when stimuli are unstructured or temporally scrambled (Dmochowski et al., 2012; Hasson, Yang, et al., 2008).

ISC is conceptually similar to measures which estimate the correlation between specific stimulus features (such as the amplitude envelope) and neural responses (cf. Ding & Simon, 2013; Iotzov & Parra, 2019; Peelle et al., 2013). However, critically, neural synchronization to stimulus features is only one potential source that may drive shared neural patterns across subjects during naturalistic listening. Engaging narratives are designed to provide a shared conscious experience to listeners — this is driven, in part, by the recruitment of processes that enable each listener to integrate incoming information with existing schemas, refine their understanding, infer what is unsaid, and anticipate what may be coming next, while filtering out distractions (Naci et al., 2014). This manifests as ongoing engagement in the story. These cognitive processes are executive in nature since they are coordinating all mental activity as the story unfolds, enabling the listener to follow along and understand. ISC is sensitive to such experiential aspects of engaging materials (Dmochowski et al., 2014; Naci et al., 2014; Nastase et al., 2019; Nummenmaa et al., 2012; Schmälzle et al., 2015; Yeshurun et al., 2017).

In the current study, we use spoken stories to investigate how challenging listening situations affect speech intelligibility and story engagement in a group of young, normal-hearing adults. We opted to measure intelligibility and ISC in separate experiments because we suspected the inclusion of the intelligibility task may influence participants’ engagement levels during story listening. Therefore, in Experiment 1, we assess intelligibility of spoken stories presented with different levels of 12-talker babble masker. In Experiment 2, we record EEG in another group of individuals while they listen to the same stories used in Experiment 1 and investigate how synchronized activity (inter-subject correlation) is affected by background masking during story listening.

## Experiment 1: Examining story intelligibility and engagement

In Experiment 1, we investigate the extent to which participants understand spoken stories when they are masked with 12-talker babble at different signal-to-noise ratios (SNRs). We further assess overall listening engagement behaviorally using a modified narrative absorption scale (Kuijpers et al., 2014) adapted for use with spoken stories (Herrmann & Johnsrude, 2020b). Speech intelligibility ratings from Experiment 1 will be related to data from Experiment 2, in which we investigate neural signatures of story engagement, to assess the extent to which story engagement relates to speech intelligibility.

## Materials and Methods

### Participants

Eighty-two individuals (mean: 28.8 years; age-range: 18-36 years; 51 males 31 females) without self-reported hearing loss, neurological or psychiatric disorders, participated in Experiment 1. All participants were recruited from the Amazon Mechanical Turk online participant pool (MTurk; https://www.mturk.com/) via the participant sourcing platform Cloud Research (previously TurkPrime; Litman et al., 2017). Participants provided informed consent and the study protocol was approved by the University of Western Ontario’s Non-medical Research Ethics Board (REB #112574). Each individual received financial compensation of $6 USD following completion of the study ($10 USD hourly rate).

Online research can be subject to increased levels of random responders as experimenters have limited control over the testing environment. However, previous work indicates that online studies generally replicate findings of in-person data collection (Berinsky et al., 2014; Buchanan & Scofield, 2018; Buhrmester et al., 2011; Gosling et al., 2004; Mason & Suri, 2012; Thomas & Clifford, 2017). Twenty-one additional individuals participated in the study but were not included, due to a technical error during data recording (N=2); reporting hearing aid usage or constant ringing in at least one ear (N=9); not wearing headphones during the study (N=3); or submitting random one-word answers to all questions on the intelligibility task (N=7).

### Acoustic stimulation and procedure

Four stories were selected from the story-telling podcast The Moth (https://themoth.org): *Reach for the Stars One Small Step at a Time* (by Richard Garriott, ∼13 min), *The Bounds of Comedy* (by Colm O’Regan, ∼10 min), *Nacho Challenge* (by Omar Qureshi, ∼11 min), and *Discussing Family Trees in School Can Be Dangerous* (by Paul Nurse, ∼10 min). Each story had 12-talker babble noise added as a masker. Babble noise was derived using the masker materials from the Revised Speech in Noise (R-SPIN) test (Bilger, 1984). Individual 5-s babble noise snippets were randomly selected from a total set of 100 and concatenated to equal the length of each story. The signal-to-noise ratio (SNR) was pseudo-randomly varied approximately every 30 to 33 s throughout each story (32, 30, 33, and 30-s, respectively, for the four stories). Five SNRs were chosen: clear, +12, +7, +2, and -3 dB signal-to-noise ratio (SNR) relative to a 12-talker babble masker. SNR was manipulated by adjusting the dB level of both the story and masker. This ensured that the overall sound level remained constant throughout a story and was similar for all stories. Three versions of SNR condition order were generated for each story in order to ensure that specific parts of a story were not confounded with a specific SNR. Within each version, SNR was varied pseudo-randomly such that a particular SNR could not be heard twice in succession. Four 30, 33, or 30-s segments per SNR were presented for stories by O’Regan, Qureshi, and Nurse, respectively, and five 32-s segments per SNR were presented for the longest story by Garriott.

For each story, phrases/sentences ranging from 4 to 8 words (range of durations: 0.62–3.6 s) were selected for intelligibility (word report) testing. These test phrases/sentences did not occur during the transition period from one SNR to the next (for approximately 5-s before, and 1-s after, the SNR transition). Four phrases per 30–33-s segment were selected, resulting in 100 phrases for the story by Garriott, and 80 for each of the other three stories. Two of the four selected phrases per 30–33-s segment were used as one intelligibility test set, whereas the other two of the four selected phrases were used as a second intelligibility test set (5 or 4 segments × 5 SNRs × 2 phrases/sentences × 2 intelligibility test sets = 100 or 80 phrases). Having two test sets ensured that the observed intelligibility effects were not confounded by specific phrases/sentences.

The experiment was conducted online, using custom written JavaScript/html and jsPsych code (Version 6.1.0, a high-level JavaScript library used for precise stimulus control; de Leeuw, 2015). The experiment code was stored at an online Gitlab repository (https://gitlab.pavlovia.org) and hosted via Pavlovia (https://pavlovia.org/). During the main experiment, each participant listened to one pseudo-randomly selected story (Garriott: N = 21; O’Regan: N = 22; Qureshi: N = 20; Nurse: N = 19). SNR order and intelligibility test set were randomly assigned. A black fixation cross was presented at the center of the screen throughout the story. The fixation cross turned yellow two seconds prior to the beginning of a test phrase/sentence, cueing the participant to prepare for intelligibility testing (see Figure 1). The fixation cross then turned green for the duration of the test phrase in the story, indicating to the participant the phrase they would be asked to report back. The story stopped with the offset of the test phrase, and an input text box appeared on the screen. Participants were asked to type their answer into the text box, after which the story resumed from the beginning of the sentence most recently heard (allowing for story continuation). After the story ended, participants answered questions that assessed comprehension and rated statements about their story listening experiences (see Table 1).

**Figure 1.**
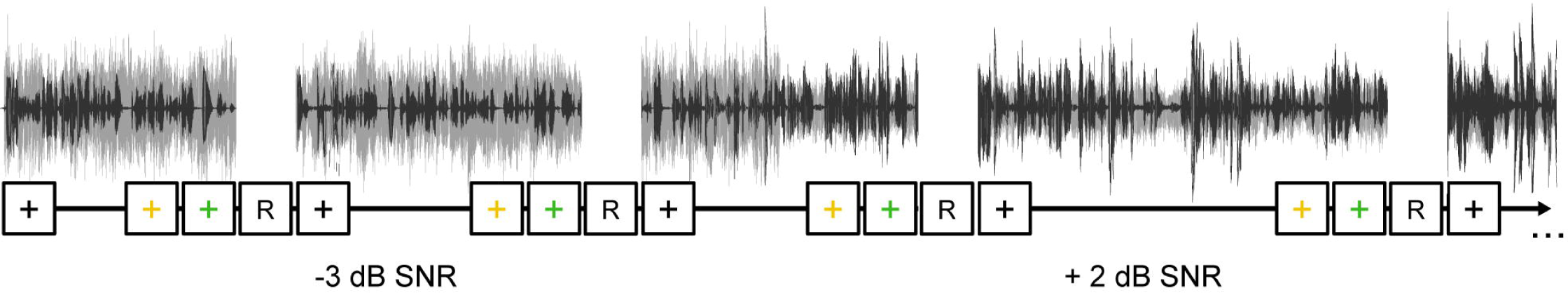
Schematic representation of the speech intelligibility task. Participants listened to a spoken story (black) masked with different levels of 12-talker babble noise (grey). A fixation cross was displayed on the computer screen throughout the study and changed colors to communicate which parts of the spoken story participants would need to recall. The fixation cross turned yellow 2-s prior to the beginning of a test phrase/sentence, cueing the participant to prepare for intelligibility testing. The fixation cross turned green during the phrase/sentence participants would be asked to report back. The story stopped with the offset of the test phrase/sentence, at which point participants would report back the phrase/sentence. The story resumed once a response was submitted.

**Table 1.** Statements of the narrative absorption scale (NAS), enjoyment, and motivation (Herrmann and Johnsrude, 2020a)

| Dimension | Statement |
| --- | --- |
| Attention (NAS) | When I finished listening I was surprised to see that time had gone by so fast. |
| Attention (NAS) | When I was listening I was focused on what happened in the story. |
| Attention (NAS) | I felt absorbed in the story. |
| Attention (NAS) | The story gripped me in such a way that I could close myself off for things that were happening around me. |
| Attention (NAS) | I was listening in such a concentrated way that I had forgotten the world around me. |
| Emotional engagement (NAS) | When I listened to the story I could imagine what it must be like to be in the shoes of the main character(s). |
| Emotional engagement (NAS) | I felt sympathy for the main character(s). |
| Emotional engagement (NAS) | I felt connected with the main character(s) of the story. |
| Emotional engagement (NAS) | I felt how the main character(s) was/were feeling. |
| Emotional engagement (NAS) | I felt for what happened in the story. |
| Mental imagery (NAS) | When I was listening to the story I had an image of the main character(s) in mind. |
| Mental imagery (NAS) | When I was listening to the story I could see the situations happening in the story being played out before my eyes. |
| Mental imagery (NAS) | I could imagine what the world in which the story took place looked like. |
| Transportation (NAS) | When I was listening to the story it sometimes seemed as if I were in the story world too. |
| Transportation (NAS) | When listening to the story there were moments in which I felt that the story world overlapped with my own world. |
| Transportation (NAS) | The world of the story sometimes felt closer to me than the world around me. |
| Transportation (NAS) | When I was finished with listening to the story it felt like I had taken a trip to the world of the story. |
| Transportation (NAS) | Because all of my attention went into the story, I sometimes felt as if I could not exist separate from the story. |
| Enjoyment | I thought it was an exciting story. |
| Enjoyment | I thought it was an enthralling story. |
| Enjoyment | I listened to the story with great interest. |
| Enjoyment | I thought the story was beautiful. |
| Enjoyment | I thought the story was presented well. |
| Motivation | I was motivated to listen to the story. |

In order to familiarize participants with the intelligibility task, a brief practice block was presented prior to the main experiment. Participants heard a ∼3-min story (a shortened version of *A Shoulder Bag to Cry On* by Laura Zimmerman), without added babble noise, and performed 12 trials of the intelligibility task (2 trials per 30-s segment).

### Online research quality assurance measures

Participants completed two initial listening tasks at the very beginning of the online session. First, participants listened to a 15-s stream of pink noise normalized to the same root-mean-square amplitude as the stories and were instructed to adjust their volume to a comfortable listening level. Participants had the option to replay the noise if they needed additional time to adjust their volume. This task ensured that participants had an opportunity to adjust their volume to a comfortable level prior to the intelligibility task, after which they were instructed to not make further adjustments.

Second, participants completed a headphone-check procedure to determine whether participants were wearing headphones as instructed (cf. Woods et al., 2017). Participants performed a tone discrimination task (6 trials; ∼2 minutes total duration), in which they determined which of three consecutive 200-Hz sine tones was the quietest. The three tones differed such that one was presented at the comfortable listening level, one at –6 dB relative to the other two tones, and one at the comfortable listening level with a 180° phase difference between the left and right headphone channels (anti-phase tone). This task is straightforward over headphones, but difficult over loudspeakers, because the pressure waves generated from an anti-phase tone interfere if heard through loudspeakers (Woods et al 2017). If they were listening through loudspeakers, they would likely falsely select the anti-phase tone as the quietest tone. No participants were excluded solely on the basis of performance on this test, however, it did provide a metric which could flag potentially non-compliant online subjects.

### Assessment of intelligibility

We calculated the proportion of correctly reported words for each SNR (Clear, +12, +7, +2, -3), across the three versions of the four stories. Different or omitted words were counted as errors, but minor misspellings and incorrect grammatical number (singular vs. plural) were not. Word-report performance was assessed using a repeated-measures analysis of variance (rmANOVA) with SNR (Clear, +12, +7, +2, -3 dB SNR) as the within-subjects factor.

### Assessment of story comprehension

Story comprehension was assessed using 8 statements, which either correctly or incorrectly described an element from the story the participant heard. Participants were asked to categorize each statement as true or false. Comprehension performance was calculated as the proportion of statements categorized correctly. Comprehension was statistically examined using a one-sample *t*-test which tested scores against chance-level performance of 0.5.

### Assessment of story engagement, enjoyment, and motivation

Following the assessment of story comprehension, we also assessed how much participants were engaged with the story, how much they enjoyed the story, and how motivated they were to listen. In order to assess story engagement, we utilized a narrative absorption scale (NAS; Kuijpers et al., 2014) adapted previously for spoken stories (Herrmann & Johnsrude, 2020b). The NAS contains statements along four dimensions (attention, emotional engagement, mental imagery, and transportation; Herrmann & Johnsrude, 2020b; Kuijpers et al., 2014) , which participants rated on a 7-point scale, where ‘1’ referred to completely disagree and ‘7’ referred to completely agree (Table 1). Participants also rated statements about enjoyment and listening motivation on the same 7-point scale (Table 1). Rating scores for each statement were averaged separately for narrative absorption (NAS) and enjoyment. Ratings for NAS and enjoyment were statistically examined in separate one-sample *t*-tests which tested ratings against a neutral response (test value: 4). Given that motivation was assessed using a single question (Table 1), the data remain ordinal rather than continuous. Motivation was therefore assessed using a one-sample Wilcoxon signed-rank test, against a neutral response (test value: 4).

### Assessment of the relationship between measures of engagement, enjoyment, and motivation

To characterize the relationship between behavioral ratings of absorption (NAS), enjoyment, motivation, and story comprehension, we calculated correlations (Pearson) between pairs of behavioral measures (6 pairs).

### Experimental design and statistical analysis

Statistical analyses were conducted using IBM SPSS Statistics for Windows (v24) and MATLAB. Details of the specific variables and statistical tests for each analysis can be found in analysis subsections for each measure. In general, effects were examined either using a rmANOVA, paired-samples *t*-tests, or one-sample *t*-tests. Behavioral ratings that were considered ordinal, not continuous, were analyzed using non-parametric tests for either one sample (Wilcoxon signed rank test) or independent sample comparisons (Wilcoxon rank sum). Significant effects were followed up using *t*-tests, with multiple comparisons corrected using the false discovery rate (FDR; Benjamini and Hochberg, 2016). FDR corrected p-values are reported as *p_FDR_.* Effect sizes are reported as partial eta squared (η^2^p ) for rmANOVAs and r_equivalent_ (r_e_ ; Rosenthal and Rubin, 2003), for *t*-tests. Greenhouse-Geisser corrected p-values are reported when sphericity assumptions have not been met (reported as *p_GG_*). This experiment was not preregistered. Data are available upon reasonable request.

## Results and Discussion

### Story comprehension, motivation, and enjoyment are high despite speech masking

Performance on comprehension questions was significantly above chance (M = .80, se = .02; *t*_81_ = 14.79, *p* = 1 × 10^-24^, r_e_ = .85), which suggests participants were able to grasp details from each story despite varying the SNR and performing the speech intelligibility task while listening to each story.

We also examined participants’ level of story engagement by testing mean ratings for NAS, enjoyment, and motivation against a rating of 4 (neutral response). Scores on the NAS were not significantly different from a neutral response (*p* = .832), but enjoyment (*t*_81_ = 2.87, *p* = .005, r_e_ = .3) and motivation (V = 2103, *p* = 5 × 10^-11^) ratings were higher than the neutral point (see Figure 2a). This suggests that although participants were ambivalent regarding whether they were fully immersed in the story they heard, they appeared to enjoy listening to the story and were motivated to do so. These findings are somewhat inconsistent with previous research on story engagement (Herrmann & Johnsrude, 2020b), which found engagement to be high and largely unaffected by moderate noise levels. The presence of the intelligibility task during story listening may have altered the listening experience such that engagement was reduced to enable detail-oriented listening to ensure participants could report back specific words when prompted.

**Figure 2.**
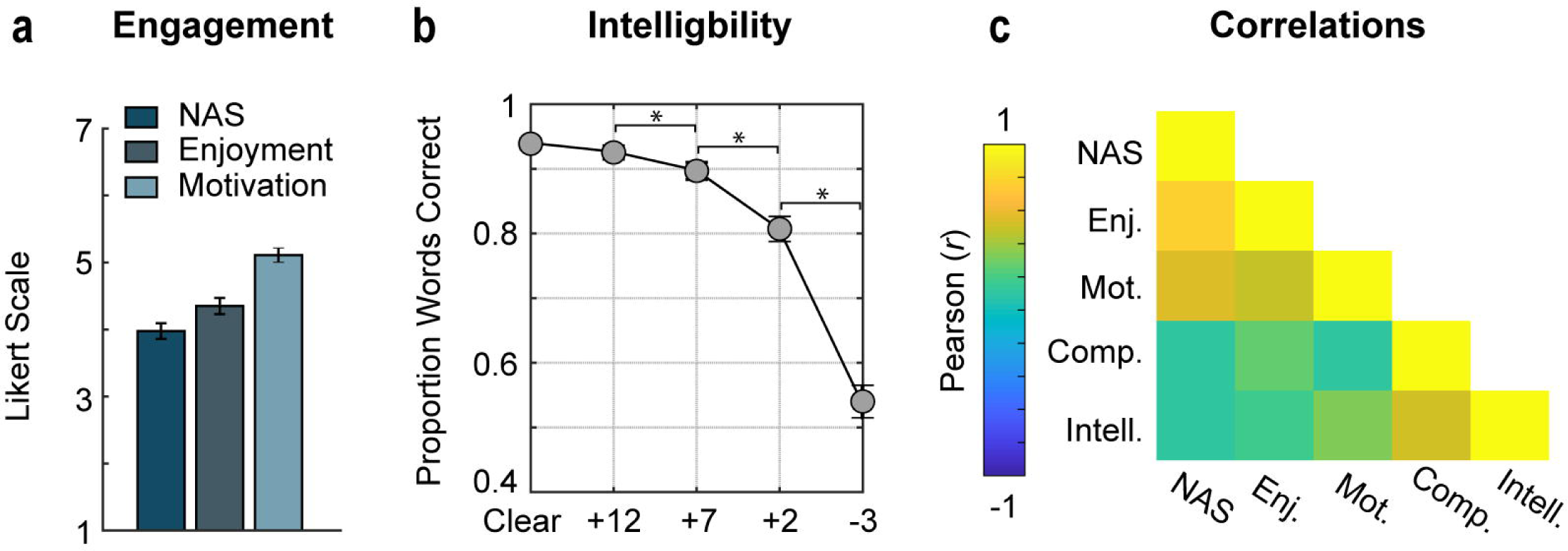
Behavioral measures of story comprehension, engagement, and intelligibility. (a) Mean ratings of the statements comprising the narrative absorption scale (NAS), and those assessing enjoyment and motivation. Ratings were provided on a 7-point Likert scale, where ‘1’ refers to completely disagree and ‘7’ refers to completely agree. (b) Mean proportion of correctly reported words plotted as a function of SNR (clear, +12, +7, +2, -3 dB). (c) Correlation matrix depicting the correlation (Pearson) between behavioral measures of story comprehension, engagement, and intelligibility. Error bars reflect the standard error of the mean. *p < 0.05.

### Story intelligibility declines as SNR decreases

For the intelligibility task, the proportion of correctly reported words declined with decreasing SNR (*F*_4,324_

= 176.5*, p* = 2.98 × 10^-38^, η^2^_p_ = .69; see Figure 2b), as predicted (Bronkhorst, 2000; Brungart, 2001; Brungart et al., 2001; Duncan & Aarts, 2006). Comparing intelligibility at successive pairs of SNRs, in order to identify the SNRs at which a significant drop in performance occurred, revealed that intelligibility was comparable between the Clear and +12 dB SNR conditions (*p_FDR_* = .077), but declined from +12 to +7 dB SNR (*t*_81_ = 3.33, *p_FDR_* = .002, r_e_ = .35), +7 to +2 dB SNR (*t*_81_ = 5.7, *p_FDR_* = 3.71 × 10^-7^, r_e_ = .54), and +2 to - 3 dB SNR (*t*_81_ =14.94, *p_FDR_* = 2.22 × 10^-24^, r_e_ = .86).

### Relating intelligibility and behavioral measures of engagement and comprehension

In an effort to quantify how intelligibility may relate to engagement and comprehension measures, we calculated correlations between intelligibility (averaged across SNRs), NAS, motivation, enjoyment, and comprehension scores (see Figure 2c). We found that intelligibility was significantly correlated with motivation (*r*_80_ = .33, *p_FDR_* = .004) and comprehension scores (*r_80_* = .50, *p_FDR_* = 6.7 ×10^-6^), such that better word-report performance was associated with higher levels of motivation to listen and better comprehension of the stories. We did not find a significant correlation between intelligibility and NAS (*r_82_* = .10, *p_FDR_* = .381) or intelligibility and enjoyment scores (*r_80_* = .16, *p_FDR_* =.223), suggesting that enjoyment/engagement with the story and being able to report the specific words spoken during the story may be independent (cf. Herrmann & Johnsrude, 2020b).

We also found a significant correlation between NAS and enjoyment (*r_80_* = .77, *p_FDR_* = 3.9 ×10^-16^), and NAS and motivation (*r_80_* = .55, *p_FDR_* = 3.4 ×10^-6^), suggesting that higher levels of engagement were associated with greater enjoyment of the stories and motivation to listen. We further observed a significant relationship between enjoyment and motivation (*r_80_* = .48, *p_FDR_* = 1.6 ×10^-5^), and enjoyment and comprehension (*r_80_* = .26, *p_FDR_* = .029). This indicates that enjoyment was associated with greater motivation to listen and better comprehension. No relationship was observed between NAS and comprehension (*r_80_* = .11, *p_FDR_* = .381), or between motivation and comprehension (*r_80_* = .12, *p_FDR_* = .367).

The results of Experiment 1 suggest that participants were able to maintain a relatively high level of story comprehension, were generally motivated, and enjoyed listening to the stories, despite the presence of the babble masker. Further, speech intelligibility was relatively good for SNRs up to +2 dB (∼80% correct words report), but substantially declined for a higher masking level (-3 dB SNR; ∼54%). The intelligibilty and engagement data obtained in this experiment will be related to inter-subject correlation during story listening in Experiment 2.

## Experiment 2: Examining inter-subject correlation during story listening

Synchronization of neural activity across different individuals – measured as inter-subject correlation – has previously been suggested to provide a window onto engagement and shared experience with naturalistic materials (Hasson et al., 2010; Nastase et al., 2019; Nguyen et al., 2019; Yeshurun et al., 2017). In Experiment 2, we investigate how inter-subject correlation during story listening is affected by 12-talker background babble noise at different SNRs.

## Materials and Methods

### Participants

Thirty-nine individuals (mean: 20.3 years; age-range: 18-32 years; 19 males 20 females) without hearing loss, neurological issues, or psychiatric disorders participated in Experiment 2. All participants were recruited from Western University or the surrounding community of London, Canada. All participants provided written informed consent and were financially compensated with $10 CAD per hour. The study was conducted in accordance with the Declaration of Helsinki, the Canadian Tri-Council Policy Statement on Ethical Conduct for Research Involving Humans (TCPS2-2014), and approved by the local Health Sciences Research Ethics Board of the University of Western Ontario (REB #112015). Five additional individuals participated but were not included either due to an issue with sound delivery (N =2), or a technical error during data recording (N=3).

### Acoustic stimulation and procedure

The experiment was conducted in a single-walled sound-attenuating booth (Eckel Industries). Sounds were delivered through Sennheiser (HD 25 Light) headphones at a comfortable listening level, using a Focusrite Scarlett 2i4 external soundcard controlled by a PC (Windows 10) and Psychtoolbox (Version 3) in MATLAB (R2017b).

Stimuli were the same four stories used Experiment 1, with the same SNRs (Clear, +12, +7, +2, -3) and the same three versions of SNR condition order per story. EEG was recorded while participants listened to each of the four stories consecutively (see Figure 3). Story version and story order were counterbalanced across participants. After each story, comprehension, story engagement, motivation, and enjoyment were assessed, and these data analyzed, as in Experiment 1. After listening to all four stories, participants completed a resting block, where they sat quietly for six minutes with their eyes open while EEG was recorded. This block was used as a baseline for ISC analyses of story listening.

**Figure 3.**
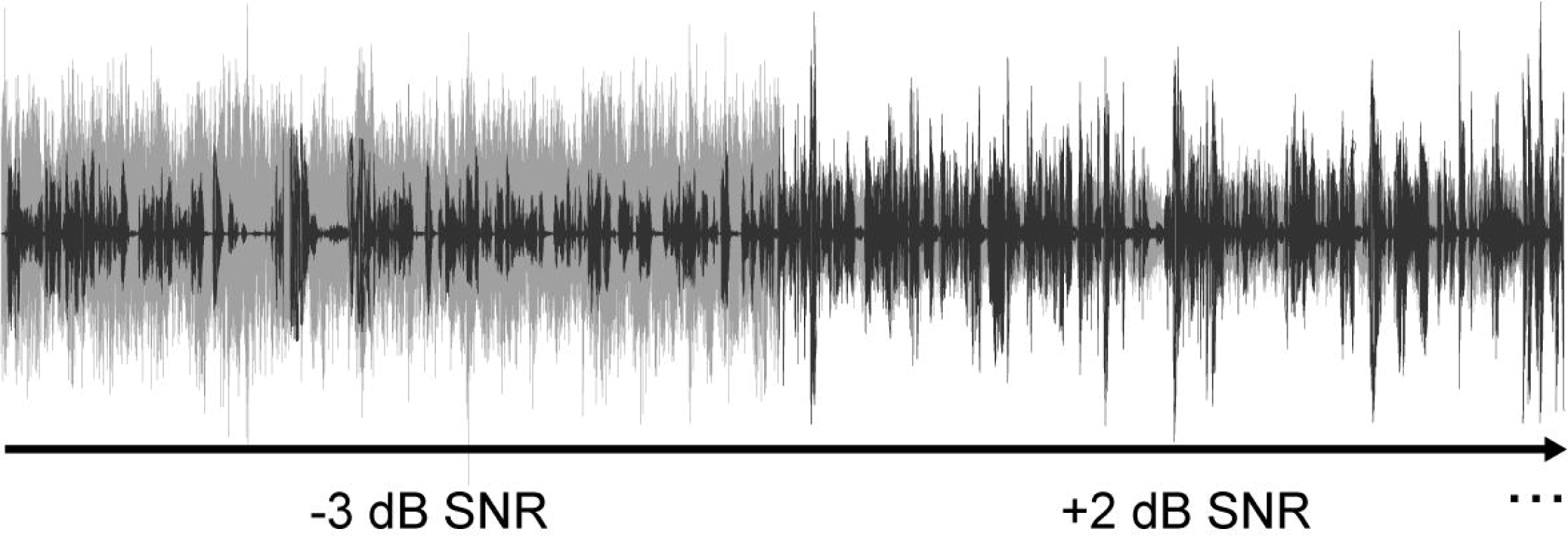
Stimulus design for Experiment 2. Participants listened to four consecutive spoken stories (black) masked with varied levels of 12-talker babble noise (grey); two of the SNRs are shown here. Each SNR lasts for ∼30-33 sec (see text for details).

### EEG recording and preprocessing

EEG was recorded from 64 active electrodes (Ag/AgCl) placed on the scalp using an electrode cap and two additional electrodes placed on both mastoids, with spacing on the scalp according to the 10/20 system (Biosemi ActiveTwo system). During data recording, all electrodes were referenced to a feedback loop of two electrodes, a common mode sense (CMS) active electrode and a driven passive electrode (see www.biosemi.com/faq/cms&drl.htm). EEG was recorded at 1024 Hz with an online low-pass filter of 208 Hz to focus on cortical sources.

All pre-processing was carried out offline using MATLAB software, the Fieldtrip toolbox (Oostenveld et al., 2011), and custom scripts. EEG data were re-referenced to the average of the signal from both mastoids. EEG data were then notch-filtered at 60 Hz to attenuate line noise, and then high-pass (0.5 Hz, 3429 points, Hann window) and low-pass filtered (22 Hz, 211 points, Kaiser window). EEG data were also downsampled to 256 Hz. Artifacts due to eye movements (saccades, blinks) and muscle activity were removed using independent components analysis (Makeig et al., 1996). To further exclude additional artifacts from subsequent analyses, data segments in which the EEG signal changed by more than 80 µV within a 0.2-s period in any channel were set to 0 µV (cf. Dmochowski et al., 2012, 2014; Cohen and Parra, 2016).

### Inter-subject correlation

We quantified inter-subject correlation (ISC) using correlated component analysis (CCA; Dmochowski et al., 2012; Parra et al., 2019), a signal decomposition method that identifies a set of electrode weights (i.e., spatial filters) yielding a linear combination of electrode activation (components) maximally correlated across participants. CCA can uncover patterns of neural activity that would not be possible with an electrode-to-electrode correlation method (S. S. Cohen & Parra, 2016; Dmochowski et al., 2012; Ki et al., 2016). For mathematical details of the method see (Parra et al., 2019), for available MATLAB scripts see (https://www.parralab.org/isc/), and for the adapted version of the MATLAB scripts used in the current study see (https://osf.io/tv7kg/). We calculated the components for each story individually to enable leave-one-out analyses (see below), such that spatial filter calculations are independent from subsequent analyses (see also Crosse et al., 2016; Herrmann et al., 2018; Broderick et al., 2019). In line with previous work, we restrict our analysis to the three components with highest overall ISC for each story (cf. Dmochowski et al., 2012, 2014; Cohen and Parra, 2016; Ki et al., 2016), because they show consistent spatial projections onto the scalp across stories (see Figure 4, left panel).

**Figure 4.**
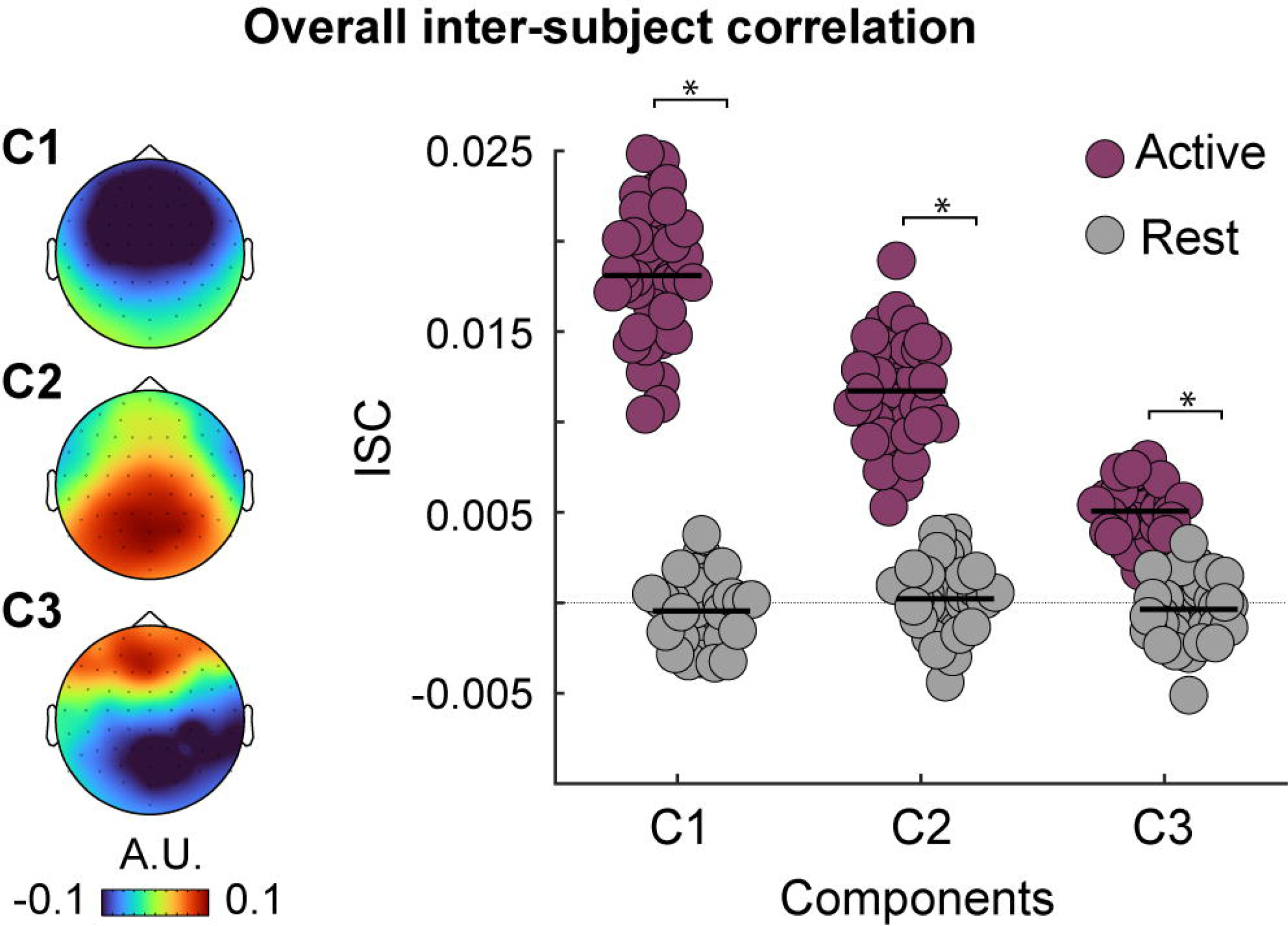
Overall inter-subject correlation. Left panel: Scalp projections of the three most correlated components. Right panel: Overall ISC is plotted as a function of component number (Component 1, Component 2, Component 3) and listening condition (active: listening to stories; rest: relaxing with eyes open). Dots represent data points for individual participants. Black solid lines indicate means. The thin, dashed line marks ISC of zero. *p < 0.05.

Separately for each of the three components (i.e., spatial filters), component weights for one story were multiplied with participants’ EEG data for the other three stories. Spatially filtered data for the three stories were then concatenated, leading to one response time course which reflects the component’s underlying sources (cf. Crosse et al., 2016; Broderick et al., 2019). This was repeated for all 4 stories, resulting in four spatially filtered EEG time courses for each component, per participant (4 concatenated datasets × 3 components = 12 spatially filtered time courses). All ISC analyses described in subsequent paragraphs were calculated after using this leave-one-out approach. ISC values were subsequently averaged across the four leave-one out iterations for each component.

For an overall ISC analysis independent of SNR conditions, each participant’s time course was correlated with the time course of each of the other participants, resulting in N-1 correlations per participant. ISC for a unique participant was calculated as the mean across these N-1 correlation values. In order to test whether story listening led to higher ISC than resting state activity in these ‘story-listening’ networks, the EEG data from the resting block were projected through each story’s spatial filters using the leave one-out approach described above. Resting ISC was calculated similarly to story-listening ISC. Mean story-listening ISC was compared to resting ISC using a paired sample t-test, separately for each of the three components. Note that results from using surrogate-data testing (Lancaster et al., 2018), by circularly shifting response time courses of participants, mirrored results from the resting-state contrasts.

To examine ISC as a function of SNR, we concatenated EEG segments with the same SNR and spatially filtered using the above leave-one-out approach. This resulted in a spatially-filtered EEG time series for each participant, SNR condition, and spatial-filter component. Separately for each SNR, each participant’s time course was correlated with the time course of each of the other participants who heard the same story version (i.e., same order of SNR conditions), and subsequently averaged across the resulting N-1 correlations for each subject. We analyzed ISC using rmANOVAs with SNR (Clear, +12, +7, +2, -3 dB) as one within-subjects factor, separately for each of the three components.

### Assessment of the relationship between inter-subject correlation and behavioral measures

To characterize the relationship between inter-subject correlation and behavioral ratings of absorption (NAS), enjoyment, motivation, and story comprehension, we calculated correlations (Pearson) between overall ISC for each of the three components and these behavioral measures.

### Comparison of ISC and speech intelligibility

In order to directly compare word-report performance in Experiment 1 with inter-subject correlation during story listening in Experiment 2, we transformed both word-report scores and ISC values to z-scores. We restricted our analysis to the component with the highest ISC (Component 1, see Figure 4, right panel). To assess the extent to which the effect of SNR differentially impacts intelligibility and ISC, we fit an exponential function to the z-scored word-report and ISC values using the following equation:

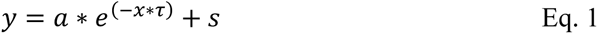

where y is the exponential function, *a* is the parameter for amplitude, *x* is a vector of linearly spaced values (1 to 5) corresponding to each SNR (clear, +12, +7, +2, -3 dB SNR), τ (tau) controls the decay strength, and the *s* parameter which allows the function to shift along the y-axis. Note that the inclusion of *s* was essential to fit the exponential function to z-scored data. To detect differences in how strongly intelligibility and ISC declined with SNR, we analyzed the τ (tau) parameter using an independent t-test, with experiment measure (intelligibility [Exp 1], ISC [Exp 2]) as the grouping variable. To make sure our results are robust to the type of function fit, we also fit a quadratic function to the z-scored word-report and ISC values, and compared quadratic and linear coefficients using two separate independent t-tests, with experiment measure (intelligibility [Exp 1], ISC [Exp 2]) as the grouping variable.

Finally, to capture how each non-clear SNR condition (+12, +7, +2, -3 dB) changed relative to the clear condition, we calculated difference scores between each SNR level and the clear condition, separately for word-report scores and ISC values. Resulting values were submitted to a mixed design ANOVA with measure (intelligibility, ISC) as the between-subjects variable and SNR (+12, +7, +2, -3 dB) as the within-subjects variable.

## Results and Discussion

### Comprehension and engagement are high despite story masking

For story comprehension, similar to Experiment 1, participants’ performance was significantly higher than chance level (M = .86, se = .01; *t*_38_ = 28.77, *p* = 2.1 × 10^-27^, r_e_ =.98), indicating that they attended and grasped details from the stories despite the varying babble masker.

We examined participants’ level of story engagement by testing mean ratings for each engagement category (NAS, enjoyment, motivation) against a rating of 4 (neutral response). In contrast to Experiment 1, scores on the NAS (*t_38_* = 3.62, *p* = .001, r_e_ = .51), enjoyment (*t_38_* =7.28, *p* = 1.02 × 10^-8^, r_e_ = .76), and motivation (*t_38_* = 8.9, *p* = 7.6 × 10^-11^, r_e_ = .82), were all significantly higher than a neutral response (see Table 2). This indicates that participants were engaged, enjoyed the stories, and felt motivated to listen. As a follow-up, we directly compared behavioral ratings between Experiment 1 and 2 using either separate independent t-tests (NAS, enjoyment) or a Wilcoxon rank sum test (motivation), with Experiment as the grouping factor (Experiment 1, Experiment 2). We found that narrative absorption (*t_119_* = 2.7, *p_FDR_* = .008, r_e_= .24), enjoyment (*t_119_* = 4.14, *p_FDR_* = 2 × 10_-4_, r_e_= .35), and motivation (Z = 2.77, *p_FDR_* = .008) were higher in Experiment 2 compared to Experiment 1 (Table 2), perhaps because Experiment 2 was in person, whereas Experiment 1 was online and Experiment 2 did not require participants to provide word reports during the story.

**Table 2.** Mean engagement (NAS), enjoyment, and motivation ratings for Experiment 1 and 2

|  | NAS | Enjoyment | Motivation |
| --- | --- | --- | --- |
| | $M \pm s.e.$ | $M \pm s.e.$ | $M \pm s.e.$ |
| Experiment 1 | $3.98 \pm 0.12$ | $4.35 \pm 0.12$ | $5.11 \pm 0.11$ |
| Experiment 2 | $4.5 \pm 0.14$ | $5.23 \pm 0.17$ | $5.59 \pm 0.18$ |

### Inter-subject correlation is higher during active listening compared to rest

Figure 4 (left panel) shows the scalp projections of the first three components (Parra et al., 2005): topographical distributions are consistent with previous work (S. S. Cohen & Parra, 2016; Dmochowski et al., 2012, 2014; Ki et al., 2016). The first component has a fronto-central scalp distribution, which is consistent with a source in auditory cortex (Näätänen & Picton, 1987; Picton et al., 2003), but may also reflect contributions from frontal cortex. The second component has a parietal scalp distribution, suggesting a source in parietal cortex or posterior auditory cortex. The third component shows a parieto-occipital peak with frontal polarity reversal.

For each component we observed ISC values that, while appearing small, were within an expected range reported previously (Dmochowski et al., 2012, 2014; Ki et al., 2016). ISC was stronger during story listening compared to rest for all three components (Component 1: *t*_38_ = 31.6, *p* = 7.3 × 10^-29^, r_e_ = .98; Component 2: *t*_38_ =19.6, *p* = 1.7 × 10^-21^, r_e_ = .95; Component 3: *t*_38_ =16.7, *p* = 4.6 × 10^-19^, r_e_ = .94; see, right panel). ISC for surrogate data (generated by circularly shifting participants’ time courses during the stories) also yielded significantly lower ISC values compared to un-shifted (story listening) data for each component. That the ISC was stronger during story listening compared to rest for the three strongest spatial components suggests that it is driven by patterns of shared neural activity evoked by the stories.

### Inter-subject correlation declines more strongly with challenging SNRs

In order to examine whether ISC is affected by SNR, we conducted an rmANOVA. Consistent with behavioral performance from Experiment 1, we observed that ISC declined with decreasing SNR for all three components (Component 1: *F_4,152_* = 20.64*, p* = 1.3 × 10^-13^, η^2^_p_ = .35; Component 2: *F_4,152_* = 18.66*, p* = 1.7 × 10^-12^, η^2^_p_ = .33; Component 3: *F* = 3.22*, p* = .014, η^2^_p_ = .08; Figure 5).

**Figure 5.**
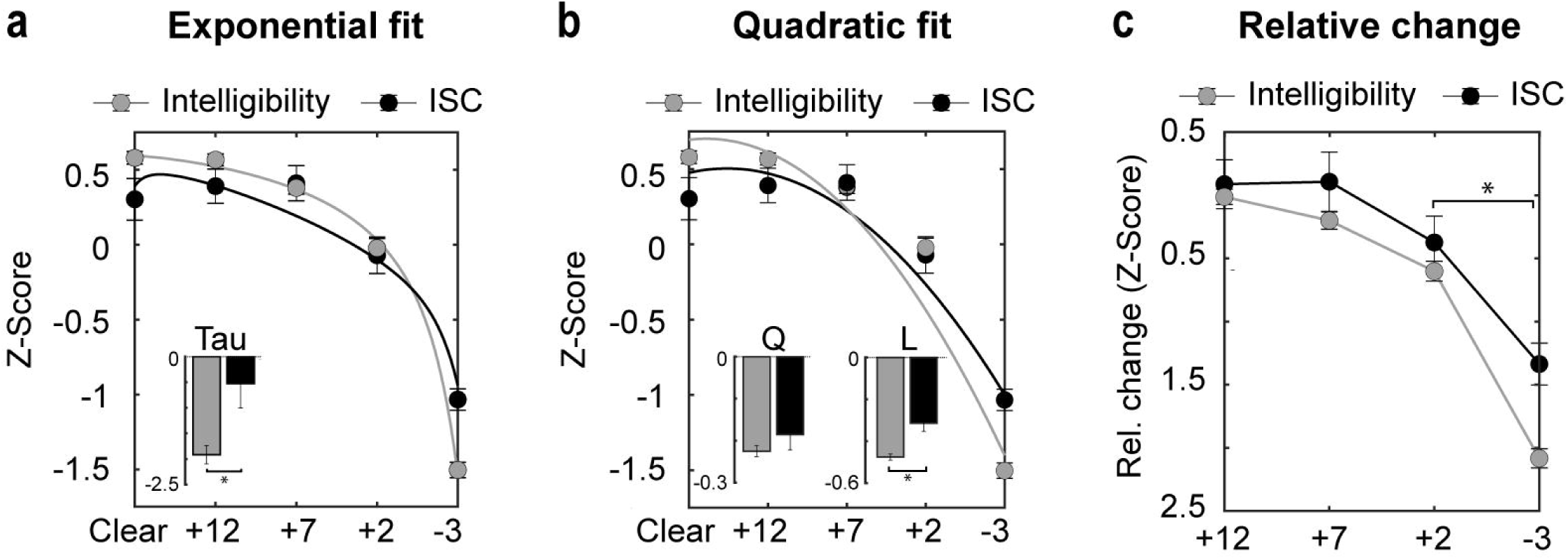
Inter-subject correlation as a function of SNR. (a) Mean ISC is plotted as a function of SNR (clear, +12,+7,+2,-3 dB) for each component (Component 1, Component 2, Component 3). Error bars reflect the standard error of the mean. *p < 0.05.

We then compared ISC at successive pairs of SNRs to identify the point at which a significant drop in ISC occurred. For Component 1, ISC strength did not differ between Clear and +12 dB SNR (*p_FDR_* = .859), or between +12 and +7 dB SNR (*p_FDR_* = .703), but decreased significantly from +7 to +2 dB SNR (*t_38_* = 2.7, *p_FDR_* = .02, r_e_ = .4), and from +2 to -3 dB SNR (*t_38_* = 5.7, *p_FDR_* = 5.2 × 10^-6^, r_e_ = .68). An identical pattern was observed for Component 2, such that ISC strength did not differ between Clear and +12 dB SNR (*p_FDR_* = .452), or between +12 and +7 dB SNR (*p_FDR_* = .498), but decreased significantly from +7 to +2 dB SNR (*t_38_* = 2.54, *p_FDR_* = .03, r_e_ = .38), and from +2 to -3 dB SNR (*t_38_* = 4.72, *p_FDR_* = .0001, r_e_ = .61). No differences were observed between successive SNRs for Component 3 after correcting for multiple comparisons (*p_FDR_ >*.14). Together, this suggests that ISC for Component 1 and 2 remained fairly stable at moderate SNRs, but decreased most substantially at the lowest SNRs. Neural ISC correlates with the degree of engagement with naturalistic materials (Hasson et al., 2010; Nastase et al., 2019; Nguyen et al., 2019; Yeshurun et al., 2017), and our data may thus suggest that listeners remained engaged during naturalistic listening even when about 10% of words were missed (+7 dB SNR; see Figure 2 for intelligibility data), but started disengaging when approximately 20% of words were missed (+2 dB SNR).

Finally, we compared the SNR condition with the lowest ISC (-3 dB SNR) to ISC during rest to determine whether the reduced synchronization observed for the most challenging listening condition was still greater than average synchronization during rest. We observed stronger ISC during the -3 dB SNR condition for Component 1 (*t_38_* = -21.44, *p_FDR_* = 2.4 × 10^-22^, r_e_ = .96), Component 2 (*t_38_* = -17.74, *p_FDR_* = 8.4 × 10^-20^, r_e_ = .94), and Component 3 (*t_38_* =-11.49, *p_FDR_* = 6.2 × 10^-14^, r_e_ =.88). Therefore, while ISC was significantly reduced at -3 dB relative to the other less challenging SNRs, observing stronger ISC at -3 dB compared to rest may suggest participants were still somewhat engaged. These results are consistent with previous behavioral work suggesting that listeners continue to engage with spoken stories despite the presence of masking and reduced speech intelligibility (Herrmann & Johnsrude, 2020b).

### Relating inter-subject correlation and behavioral measures of engagement and comprehension

Correlations between overall ISC for each component (Component 1, Component 2, Component 3) and behavioral ratings of absorption (NAS), enjoyment, motivation, and story comprehension were not significant after correcting for multiple comparisons (*r_37_* < .3, *p_FDR_* > .22).

Consistent with Experiment 1, we found a significant correlation between NAS and enjoyment (*r_37_* = .81, *p_FDR_* = 6.6 × 10^-9^), and NAS and motivation (*r_37_* = .70, *p_FDR_* = 4 × 10^-6^). Higher levels of absorption were associated with greater enjoyment of the stories and motivation to listen. We also found a significant relationship between enjoyment and motivation (*r_37_* = .86, *p_FDR_* = 5.8 × 10^-11^): enjoyment was associated with greater motivation to listen. NAS did not correlate with story comprehension (*r_37_* =.21, *p_FDR_* = .192), and no other significant correlations were found (*r_37_* <.23, *p_FDR_* >.36).

### Assessing the relative effect of SNR on intelligibility and inter-subject correlation

We directly compared the effect of SNR on intelligibility and inter-subject correlation by fitting an exponential function z-scored word-report values and z-scored ISC values for the component with the strongest ISC (Component 1), and compared the resulting decay coefficients between word report and ISC. We observed a larger τ coefficient for the intelligibility task relative to ISC (*t_119_* = -21.44, *p* = .001, r_e_ = .29), suggesting that speech intelligibility declined more strongly with SNR than ISC (Figure 6a). To confirm that our results are robust to the type of function used to describe the data, we also fit a quadratic function and compared the resulting quadratic and linear coefficients between word report and ISC (Figure 6b). The quadratic coefficients did not differ between word report and ISC (*p* = .21). However, we observed more negative linear coefficients for the intelligibility task relative to ISC (*t*_119_ = 4.71, p = 7 × 10^-6^, r_e_ = .4), further confirming that speech intelligibility declined more strongly with SNR than ISC.

**Figure 6.**
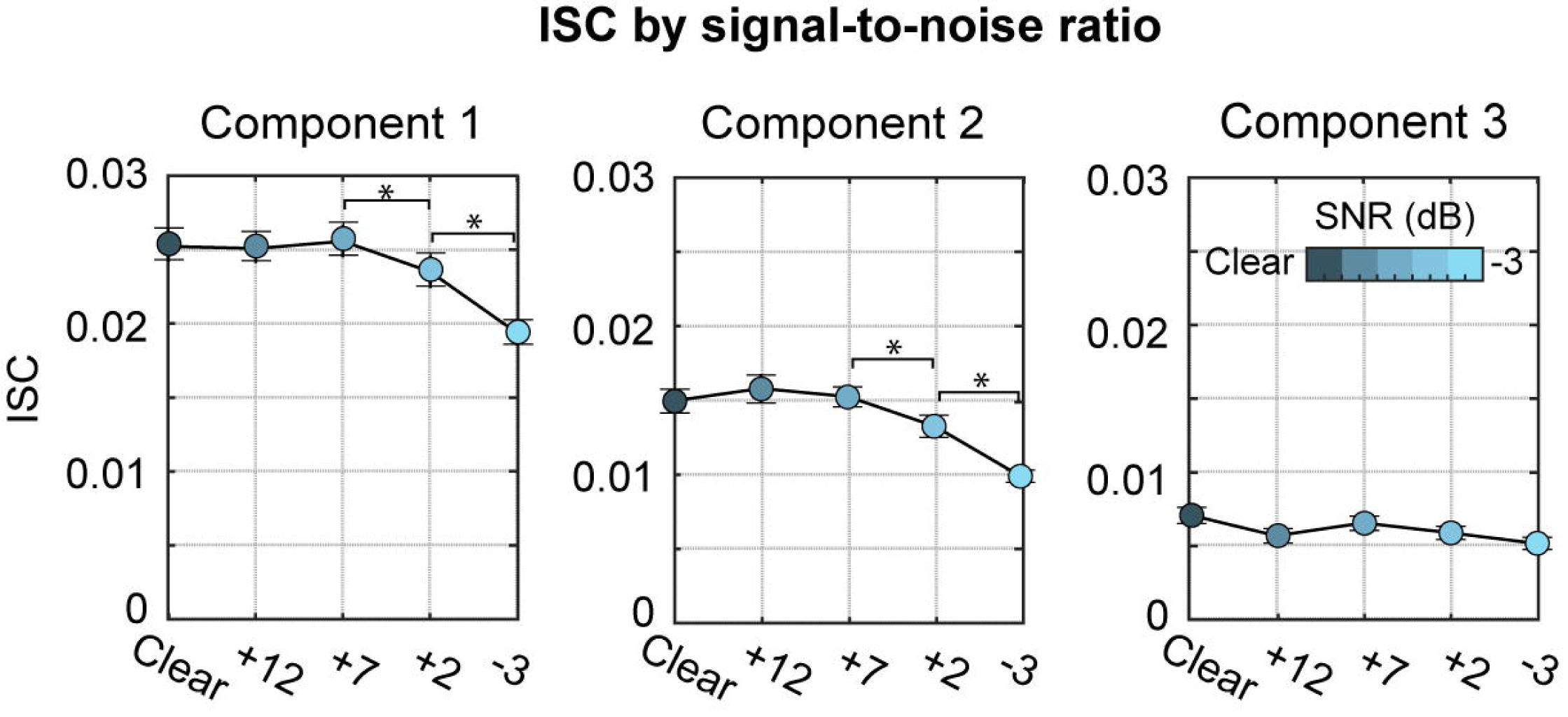
Normalized measures of intelligibility and inter-subject correlation. (a) Mean normalized word report (Exp 1) and ISC (Exp 2: Component 1) are plotted as a function of SNR (clear, +12, +7, +2, - 3 dB). Solid lines correspond to an exponential function fit to normalized means. Mean tau coefficients (decay strength) are plotted as a function of experiment measure (intelligibility, ISC). (b) Mean normalized word report (Exp 1) and ISC (Exp 2: Component 1) are replotted as a function of SNR (clear, +12, +7, +2, -3 dB). Solid lines correspond to a quadratic function fit to normalized means. Mean quadratic (left plot) and linear coefficients (right plot) are plotted as a function of experiment measure (intelligibility, ISC). (c) The change in normalized word-report (z-score) and ISC for each SNR relative to clear. Error bars reflect standard error of the mean. *p < 0.05.

In order to capture at which SNR intelligibility and ISC diverged, we calculated difference scores, separately for mean-normalized word-report and ISC, between each SNR and the clear condition and analyzed the resulting z-scores using an ANOVA. The relative change from clear for both word-report and ISC decreased significantly as SNR decreased (effect of SNR: *F*_3,357_ = 197.5, *p* = 1.8 × 10^-75^), but the decline from 2 to -3 dB SNR was larger for word report than for ISC (Task × SNR interaction: *F*_3,357_ = 5.99, *p* = .001, η^2^_p_ = .05; WS contrast: *F* =7.76, *p*_FDR_ = .01, η^2^_p_ = .06; see Figure 6c). Insert Figure 6

## General Discussion

We investigated how acoustic masking affects speech intelligibility, engagement, and neural activity during story listening. In Experiment 1, we observed that speech intelligibility declined with decreasing SNR, such that intelligibility was relatively high (80% or higher) at all but the lowest SNR (-3 dB), at which word report accuracy declined to 54%. In Experiment 2, we recorded EEG while individuals listened to spoken stories and investigated inter-subject correlation (ISC) as a possible window into engagement. We observed that ISC remained stable across all but the lowest SNRs, whereas speech intelligibility measured in Experiment 1 declined more rapidly with decreasing SNR. Our analyses suggest that speech intelligibility is less robust to changes in SNR than synchronized neural activity (ISC). Speech intelligibility may thus be an insufficient predictor of naturalistic speech listening. Our study suggests that individuals may continue to engage with naturalistic, spoken stories even under adverse listening conditions that reduce intelligibility by 10%.

### Story comprehension and overall engagement remain high despite background noise

In both experiments, participants scored well above chance on the comprehension questions, indicating that they were able to encode details from the stories despite regular changes in the level of background noise. Participants’ self-rated story absorption in Experiment 1 (our behavioral measure of engagement) was neutral, but they reported being motivated and enjoyed listening to the stories. In Experiment 2, participants reported increased levels of absorption, enjoyment, and motivation relative to neutral, which is consistent with previous research suggesting engagement and motivation for interesting narratives are largely resistant to moderate levels of noise (Herrmann & Johnsrude, 2020b). In addition, participants in Experiment 2 reported higher levels of absorption, enjoyment, and motivation relative to Experiment 1 (see Table 2).

The differences in behavioral engagement metrics between experiments may have several explanations. Engagement with narrative materials requires individuals to generate and maintain mental models of events, situations, and characters (Mar & Oatley, 2008; Oatley, 2016; Zwaan, 2016; Zwaan et al., 1995). Disruptions to mental model generation and maintenance may reduce engagement with a narrative (Busselle & Bilandzic, 2009). In Experiment 1, stories were briefly discontinued about every 10-20 seconds and participants performed the intelligibility task, whereas stories were played without interruption in Experiment 2. Moreover, Experiment 1 was conducted online, whereas Experiment 2 was conducted in-person in the lab, with possibly fewer distractions for participants. Generation and maintenance of a mental model may thus have been more disrupted in Experiment 1 than 2, leading to lower narrative absorption scores in Experiment 1 (Table 2). Alternatively, the different levels of reported engagement between the two experiments may reflect differences in listening strategy. The word-report task in Experiment 1 may have led listeners to adopt a detail-oriented strategy to ensure they could report back specific words when prompted. In contrast, the absence of a dual task in Experiment 2 may have motivated a ‘gist’ or ‘global’ strategy (Harding et al., 2007), in which individuals focus on understanding the message but are not remembering words verbatim. The conditions of Experiment 2, in which people listened without a secondary intelligibility task, is more similar to real life, and suggests that, consistent with previous work (Herrmann & Johnsrude, 2020b), engagement with spoken stories – which likely relies on gist perception – is unaffected even if a listener misses 10% of words.

### Speech intelligibility declines with decreasing signal to noise ratio

We utilized spoken stories masked with varied levels of background noise to approximate naturalistic listening conditions and found that speech intelligibility decreased as the SNR decreased. This is consistent with previous studies investigating how speech intelligibility changes with increasing speech degradation or with masking by short speech utterances, such as disconnected sentences (Akeroyd, 2008; Davis & Johnsrude, 2003; Duncan & Aarts, 2006; Obleser et al., 2007; Wild, Davis, et al., 2012; Wild, Yusuf, et al., 2012). This is also consistent with other studies using indirect measures of speech intelligibility during naturalistic listening, such as using comprehension or fill-in-the-blank questions (Best et al., 2016; Humes & Dubno, 2010; MacPherson & Akeroyd, 2013; Power et al., 2012; Xia et al., 2017). One critical difference is that our paradigm directly measures intelligibility by asking participants to report back phrases just after being heard, which allows for a more direct comparison with previous psychoacoustic work using isolated speech utterances.

We further demonstrate that intelligibility was relatively high at moderate SNRs (>80% words reported for +2 dB SNR and higher) but was particularly poor when the SNR was low (∼55% words reported for -3 dB SNR). Performance at moderate SNRs may have benefited from the influence of contextual information, as semantic or lexical context can facilitate intelligibility when the speech signal is degraded or an individual has impaired hearing (Desjardins & Doherty, 2014; Dubno et al., 2000; Holmes et al., 2018; Obleser et al., 2007; Pichora-Fuller et al., 1995). This strategy was perhaps not possible for the lowest SNR, at which only about half of the words could be understood – the context gleaned from this proportion of sentences may not be sufficient to enhance comprehension.

### Inter-subject correlation is stronger when listening to stories compared to rest

Inter-subject correlation (ISC) captures the aspects of neural activity that are synchronized across participants, typically while they watch a movie or listen to a spoken story (Dmochowski et al., 2012, 2014; Hasson et al., 2004; Hasson, Furman, et al., 2008). We observed that ISC was stronger during story listening compared to rest for all three spatial components. This is consistent with previous studies showing that ISC is stronger when attending to a coherent narrative compared to when attention is directed elsewhere (S. S. Cohen et al., 2018; Ki et al., 2016; Kuhlen et al., 2012; Rosenkranz et al., 2021) or compared to rest (Hasson et al., 2004; Wilson et al., 2008). Inter-subject correlation is often thought of as a neural signature of engagement, because ISC is stronger for captivating or exciting stimuli (Hasson et al., 2010; Schmälzle et al., 2015) and predicts behavioral engagement measures (S. S. Cohen et al., 2017; Dikker et al., 2017; Dmochowski et al., 2014; Poulsen et al., 2017; Song et al., 2021). Observing significant ISC despite the presence of background noise in the current study may suggest that participants were able to remain engaged despite missing bits of the story during periods when the SNR was low.

Critically, the term engagement is not always used in the same manner. In literature and media studies, engagement is typically defined as a multidimensional construct that includes attention, emotional engagement, mental imagery, and “transportation”, the experience of being absorbed in the world of the story (Busselle & Bilandzic, 2008, 2009; J. Cohen, 2001; Green et al., 2004; Herrmann & Johnsrude, 2020b; Kuijpers et al., 2014; see Table 1). Engagement measured as synchronized neural activity during movie watching or story listening is thought to be driven by a shared experience across individuals (Dmochowski et al., 2012; Hasson et al., 2010; Nastase et al., 2019; Nguyen et al., 2019; Yeshurun et al., 2017) and thus may converge with more experiential, cognitive engagement dimensions. Our finding that activity is less synchronized across participants when they rest and their minds are not focused on a story (Figure 4) may suggest that EEG-recorded ISC is perhaps sensitive to one or more of the aforementioned experiential dimensions of engagement. However, this remains to be more directly addressed in future research.

**Inter-subject correlation is more robust to changes in signal to noise ratio than speech intelligibility** We observed that ISC was reduced for speech listening during challenging SNRs, but not for moderate SNRs. Specifically, ISC did not differ between clear, +12, and +7 dB SNR, despite our separate behavioral experiment indicating participants missed approximately 10% of words for +7 dB SNR (Figure 2). However, ISC significantly declined from +7 to +2 and from +2 dB to -3 dB SNR. Despite similar overall patterns, comparing ISC and intelligibility directly demonstrated that word-report performance declined more strongly with decreasing SNR compared to ISC (Figures 6a - 6c). This is consistent with related work that investigated the relation between speech intelligibility and neural synchronization with the amplitude envelope of speech. Synchronization with the speech envelope appears robust to moderate changes in SNR or the presence of a competing speech stream, while intelligibility declines (Ding & Simon, 2012, 2013). Additionally, while ISC was lowest at -3 dB SNR, ISC for all three components was still stronger than during rest, which may suggest participants were still somewhat engaged when speech intelligibility was ∼55%. This is consistent with a recent behavioral study demonstrating that listeners were as absorbed during a story that was continuously masked by babble at +4 dB SNR as they were during a clear story, despite much higher rated listening effort for the former (Herrmann & Johnsrude, 2020b). The current data provide neural evidence that across-participant synchronization during story listening is unaffected by masking that reduces intelligibility to about 90%, but begins to decrease when intelligibility reduces to 80%. We speculate that observing synchronized activity despite acoustic masking may have resulted from participants accurately perceiving the story gist, and enjoying it, which perpetuated their intrinsic motivation to continue listening and stay engaged (Eckert et al., 2016; Herrmann & Johnsrude, 2020a; Matthen, 2016; Richter, 2016).

Listening and trying to understand masked speech is supported by processes such as sustained and selective attention (Davis & Johnsrude, 2007; Obleser et al., 2007; Wild, Yusuf, et al., 2012) which, when recruited, result in effortful listening (Pichora-Fuller et al., 2016). When the SNR is so poor that the gist is lost, or attentional resources become depleted (Hornsby et al., 2016; Ivarsson & Arlinger, 1994), this may lead to temporary disengagement or attentional lapses (Hallberg & Carlsson, 1991; Heffernan et al., 2016). This is consistent with our observation of lowest, but still significant, ISC at the most difficult SNR (-3 dB SNR) for which participants missed about ∼45% of words on average. Our study suggests that a listener can engage and follow a story’s thread as long as intelligibility is over 80%. However, one limitation of the current study is that intelligibility and EEG-measured ISC were assessed in separate groups of participants. Future studies may consider using a within-subjects design to more directly assess how intelligibility relates to synchronized activity across participants.

### Relating inter-subject correlation with behavioral measures of engagement

To assess the relationship between the behavioral and neural measures of engagement we calculated correlations between the overall ISC for each component (Component 1, Component 2, Component 3) and behavioral ratings of absorption (NAS), enjoyment, and motivation, but did not observe any significant relationships after correcting for multiple statistical comparisons (*r_37_* < .3, *p_FDR_* > .22). This is inconsistent with previous literature which demonstrates that ISC strength is predictive of engagement (S. S. Cohen et al., 2017; Dikker et al., 2017; Dmochowski et al., 2014; Poulsen et al., 2017; Song et al., 2021) as well as other measures dependent on sustained attention, such as subsequent recall of the materials (Chan et al., 2019; S. S. Cohen et al., 2018; S. S. Cohen & Parra, 2016; Davidesco et al., 2019; Hasson, Furman, et al., 2008; Piazza et al., 2021; Song et al., 2021; Stephens et al., 2010). We speculate that the behavioral measure of engagement did not correlate with ISC because we did not continuously assess changes in behavioral engagement throughout each story, as has been done previously (S. S. Cohen et al., 2017; Dmochowski et al., 2014; Song et al., 2021). Post-hoc measures of behavioral engagement (as we used here) have also been used on occasion: for example, relating ISC with engagement in the classroom (Dikker et al., 2017; Poulsen et al., 2017), but the listening conditions did not vary systematically, as in the current study. We changed SNR every 30 seconds - this may have led to dynamic changes in engagement (especially at high masking levels), which would be reflected in ISC, but which would be difficult to capture behaviorally with a single post-hoc measure. Further, in contrast to previous work, we used a story absorption scale (Herrmann & Johnsrude, 2020b; Kuijpers et al., 2014) as a behavioral engagement measure, but whether this accurately reflects neural engagement is a matter for further research. It would be helpful to combine our story paradigm with a continuous measure of behavioral engagement to further explore this topic (cf. Song et al., 2021).

### Using narratives to approximate realistic listening scenarios

Everyday listening situations involve speech material comprised of sentences that relate to each other, and that are interesting to the listener. Such listening situations are commonly subject to increased levels of background noise, but are also rich with many positive aspects of listening that may drive motivation to listen (Herrmann & Johnsrude, 2020a; Matthen, 2016). Similarly, the spoken stories used here are intrinsically motivating to a listener, reflecting such everyday listening situations (Dunlop & Walker, 2013). Our work shows, in line with recent studies (Brodbeck et al., 2020; Broderick et al., 2018, 2019, 2020; Erb et al., 2020; Polonenko & Maddox, 2021), that utilizing naturalistic, spoken stories to investigate speech listening provide a useful avenue to investigate listening in ecologically valid conditions.

## Conclusions

We utilized naturalistic spoken stories to investigate how challenging listening affect intelligibility and listener engagement. We used inter-subject correlation of electroencephalographic activity as a measure of listener engagement. Inter-subject correlation was unaffected by moderately challenging SNRs, despite speech-intelligibility scores suggesting listeners missed approximately 10% of words. Our data further show that speech intelligibility declines faster than electrophysiologically measured listening engagement under masked speech conditions. Our work provides a potentially fruitful approach with naturalistic, spoken stories to investigate intelligibility and engagement during masked speech listening. Our approach may importantly open future avenues to investigate (dis)engagement in older adults with hearing impairment.

## Acknowledgements

This research was supported by the Canadian Institutes of Health Research (MOP133450 to I.S. Johnsrude). BH was supported by a BrainsCAN Tier I postdoctoral fellowship (Canada First Research Excellence Fund; CFREF) and the Canada Research Chair program.

